# Anatomical Structures, Cell Types, and Biomarkers Tables Plus 3D Reference Organs in Support of a Human Reference Atlas

**DOI:** 10.1101/2021.05.31.446440

**Authors:** Katy Börner, Sarah A. Teichmann, Ellen M. Quardokus, James Gee, Kristen Browne, David Osumi-Sutherland, Bruce W. Herr, Andreas Bueckle, Hrishikesh Paul, Muzlifah A. Haniffa, Laura Jardine, Amy Bernard, Song-Lin Ding, Jeremy A. Miller, Shin Lin, Marc Halushka, Avinash Boppana, Teri A. Longacre, John Hickey, Yiing Lin, M. Todd Valerius, Yongqun He, Gloria Pryhuber, Xin Sun, Marda Jorgensen, Andrea J. Radtke, Clive Wasserfall, Fiona Ginty, Jonhan Ho, Joel Sunshine, Rebecca T. Beuschel, Maigan Brusko, Sujin Lee, Rajeev Malhotra, Sanjay Jain, Griffin Weber

## Abstract

This paper reviews efforts across 16 international consortia to construct human anatomical structures, cell types, and biomarkers (ASCT+B) tables and three-dimensional reference organs in support of a Human Reference Atlas. We detail the ontological descriptions and spatial three-dimensional anatomical representations together with user interfaces that support the registration and exploration of human tissue data. Four use cases are presented to demonstrate the utility of ASCT+B tables for advancing biomedical research and improving health.

## 2. Main

Launched thirty years ago at a cost of $3 billion, the international Human Genome Project achieved the remarkable feat of determining the sequence of base pairs in human DNA and creating a map of all the genes in the human genome^1, 2^. Decoding the human genome transformed our collective understanding of human biology and has had innumerable benefits for medicine, including drug development, disease diagnosis, and cancer treatment, among others.

There are several ongoing ambitious efforts to map the cells of the human body, the functional outcome of the transcribed genome. These initiatives aim to create a digital reference atlas of the entire healthy adult human body—from the three-dimensional shape of whole organs and thousands of anatomical structures down to the interdependencies between trillions of cells and the biomarkers that characterize and distinguish different cell types. The goal is to establish a benchmark reference that will help us understand how the healthy human body works and changes that occur when disease strikes. The consortia coordinating this work have brought together individuals from around the globe and from across disciplines and funding boundaries with expertise in high-resolution and high-throughput technologies. **Fig. 1a** shows a bimodal network of 16 consortia that focus on human atlas construction and the 30 organs they study, including efforts funded by the National Institutes of Health (NIH, blue) and cross-fertilized by the international Human Cell Atlas (HCA, red)^3, 4^. Consortia include the Allen Brain Atlas^5^, the Brain Research through Advancing Innovative Neurotechnologies Initiative - Cell Census Network (BICCN) Initiative^6^, Chan Zuckerberg Initiative Seed Networks for HCA (CZI)^3, 4, 7^, HCA awards by EU’s Horizon 2020 program, the Genotype-Tissue Expression (GTEx) project^8^, the GenitoUrinary Developmental Molecular Anatomy Project (GUDMAP)^9^, Helmsley Charitable Trust: Gut Cell Atlas^3, 4, 7^, Human Tumor Atlas Network (HTAN)^10^, Human Biomolecular Atlas Program (HuBMAP)^11^, Kidney Precision Medicine Project [KPMP]^12^, LungMAP^13^, HCA grants from the United Kingdom Research and Innovation Medical Research Council (MRC)^14^, (Re)building the Kidney (RBK)^15^, Stimulating Peripheral Activity to Relieve Conditions (SPARC)^16^, The Cancer Genome Atlas (TGCA)^17–19^, and the Wellcome funding for HCA pilot projects HCA^3, 4, 7^. In total, over 2000 experts from around the globe are working together to construct an open, digital human reference atlas using a wide variety of single-cell methods, ranging from proteomics to antibody-based imaging methods as well as single-cell genomics and imaging.

**Fig. 1.**
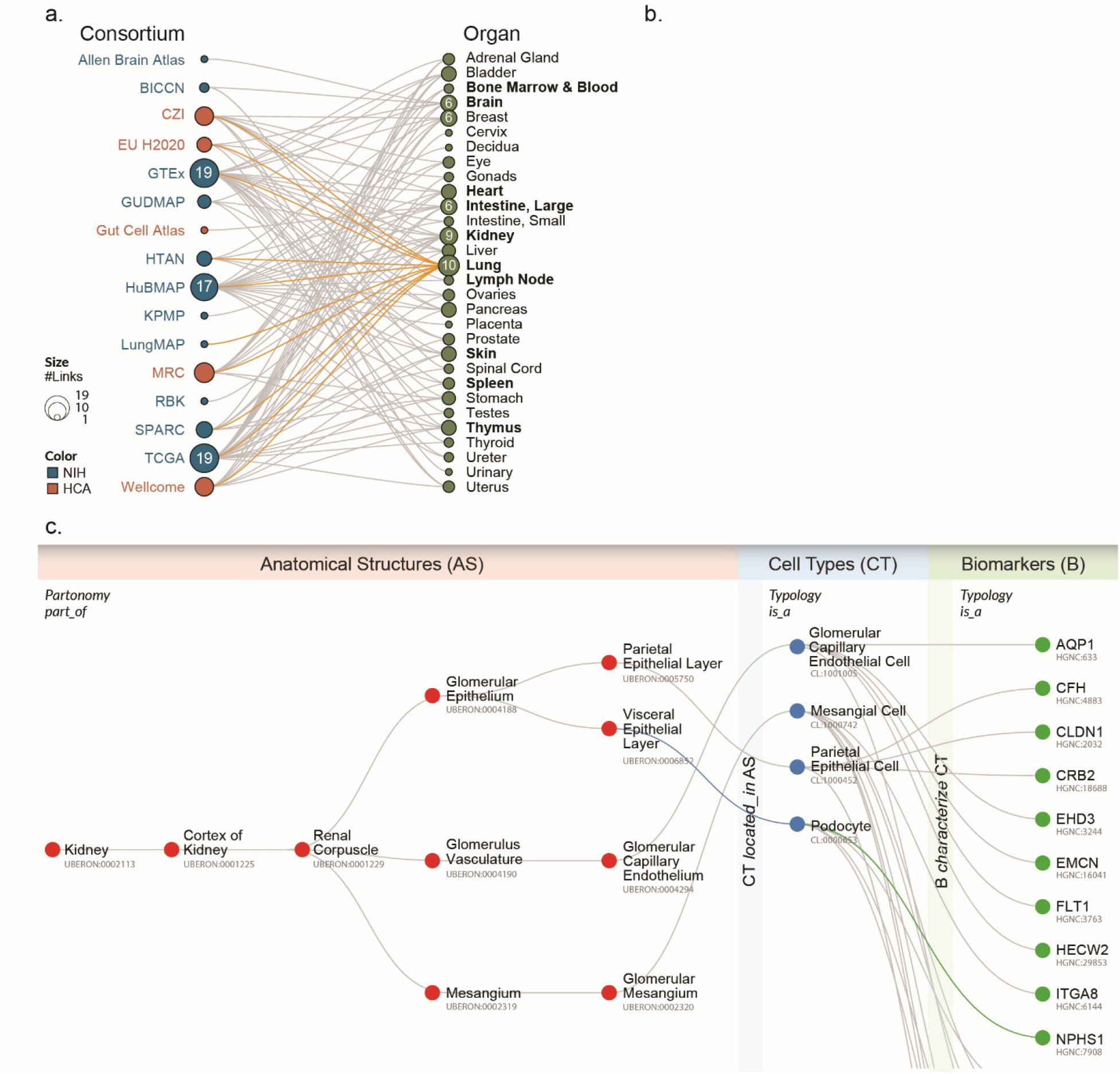
**a**, Alphabetical listing of 16 human reference atlas construction efforts (left) linked to 30 human organs they study (right). See legend for color and size coding. The lung is studied by 10 consortia; see orange links. This review focuses on 10 organs (bolded) plus vasculature. **b**, 3D reference objects for major anatomical structures were jointly developed for 11 organs. **c**, Exemplary ASCT+B table showing all AS and CT and some B for the glomerulus in kidney; annotated with names of the three entity types (AS, CT, B) and four relationship types (in italics).

A major challenge in constructing a human reference atlas is combining the data generated by the different consortia without a common “language” shared across them for describing and indexing the data. For example, different cell types can be assigned using existing ontologies/nomenclatures based on genetic, protein, or other biomarker expression profiles. Rapid progress on single-cell technologies has led to an explosion of cell-type definitions, but no standards exist for the naming of anatomical structures, cell types, and biomarkers across organ systems (but see^20^). Furthermore, information on what cell types are commonly found in which anatomical structures and what biomarkers best characterize certain cell types is scattered across hundreds of ontologies (e.g., Uberon multi-species anatomy ontology, Foundational Model of Anatomy Ontology [FMA], Cell Ontology [CL], or HUGO Gene Nomenclature) and thousands of publications (e.g., see atlas efforts for brain^21^, heart^22^, lung^22^, and kidney^12^) on cells identified during human development, disease, and across multiple species. Some critically important details (e.g., shape and distribution of microanatomical structures or the spatial layout of functionally interdependent cell types) are captured via hand-drawn figures—not digitally. This state of affairs impedes progress in biomedical science and practice as data is difficult or impossible to manage, compare, harmonize, and use.

To address these issues, an NIH-HCA organized meeting in March 2020, brought together leading experts to agree upon major ontologies and associated 3D anatomical reference objects and to expand them as needed to capture the healthy human adult body. Over the last 10 months, more than 50 experts—including physicians, surgeons, anatomists, pathologists, experimentalists, and representatives from the various consortia—have agreed on a systematic digital representation of relevant knowledge that can be used to integrate and analyze massive amounts of heterogeneous data. Specifically, experts agreed on a data format and major ontologies to be used to create a ‘Rosetta Stone’ across existing anatomy, cell, and biomarker ontologies. As a proof of concept, experts compiled inventories for 11 major organs (bolded in **Fig. 1a** and shown in **1b**)—listing known anatomical structures (AS), the cell types (CT) they contain, and the biomarkers (B) that are commonly used to characterize CT (e.g., gene, protein, lipid, or metabolic markers). The 11 organs include: bone marrow and blood plus pelvis reference organ^23–36^, brain^5, 20, 21, 37, 38^, heart^39, 40^, large intestine^41–52^, kidney^12, 53–61^, lung^62–71^, lymph node^72–82^, skin^83–95^, spleen^74, 96–107^, thymus^108–119^, and vasculature^62, 120–126^. Because spatial position and context matters for cell function, the experts collaborated with medical designers to compile 3D, semantically annotated reference objects that cover the anatomically correct size and shape of major AS in a systematic and computable manner (see final set in **Fig. 1b)**. Results are captured in so-called ASCT+B tables that list and interlink AS, CT, and B entities, their relationships, as well as references to supporting publications.

The initial tables and reference objects are not complete, but they demonstrate how existing knowledge can be captured and reorganized in support of a human reference atlas. Entities and relationships in the tables are linked to major ontologies, and existing source publications are referenced. The tables capture data and expertise that is mandatory for compiling a comprehensive human reference atlas, and they are an important tool for facilitating collaboration among the 16 consortia. Specifically, the tables and associated 3D reference objects: (1) provide an agreed-upon framework for experimental data annotation across organs and scales (i.e., from whole body to organs, tissues, cell types, and biomarkers); (2) make it possible to compare and integrate data from different assay types (e.g., scRNAseq and MERFISH data) for spatially equivalent tissue samples; (3) are a semantically and spatially explicit reference for “healthy” tissue and cell identity data that can then be compared against disease settings; and (4) can be used to evaluate progress on the semantic naming and definition of cell types and their spatial characterization. Importantly, they help communicate data structures and user needs to programmers in support of user interfaces that support the construction and usage of a human reference atlas.

The remainder of this paper details the data format, design, and usage of ASCT+B tables and associated 3D reference objects. It presents ten initial ASCT+B tables interlinked via a vasculature table together with a reference library of major AS. We then discuss four concrete use cases that showcase the utility of the initial 11-organ human reference atlas for tissue registration and exploration, data integration, understanding disease, and measuring progress toward a more complete human reference atlas. We conclude with a discussion of next steps and an invitation to collaborate on the construction and usage of a reference atlas for healthy human adults.

## 3. ASCT+B Tables

In 2019, the Kidney Precision Medicine Project (KPMP) project published a first version of the ASCT+B tables to serve as a guide to annotate structures and cell types across multiple technologies to appreciate cellular diversity in the kidney^12^. At their core, the tables represent three entity types (AS, CT, B) and five relationship types (in bold italics in **Fig. 1c**). AS are connected via *part_of* relationships creating a partonomy tree, CTs are linked via *is_a* relationships (e.g., T cell is an immune cell), and biomarkers can be of different types indicated by *is_a* (e.g., gene, protein, lipid, metabolite). Two bimodal networks link CT and AS (i.e., one and the same CT might be *located_in* multiple AS, while a single AS may have multiple CT) and CT and B (i.e., one and the same B might be used to *characterize* different CT, and multiple B might be required to uniquely *characterize* one CT).

The current ASCT+B v1.0 table format captures the AS partonomy (unlimited number of levels), one level of CT (but see extensive CT typology discussed in^127^), and two B types: gene markers (BG) and protein markers (BP). Each row in the table represents one CT *located_in* a specific AS together with all B commonly used to identify this CT. In addition, the tables include citations to relevant work documenting not only AS, CT, and B and their interlinkages, but also AS-CT and CT-B relationships.

The initial set of 11 tables was authored using templated Google sheets. For each AS, CT, and B entity, authors completed (1) its preferred name, (2) an ontology name/label, if available, and (3) a unique, universally resolvable ontology ID, if available. A lookup table of organ-specific human, non-developmental data captured in existing formalized ontologies (initially Uberon for AS, Cell Ontology [CL] for CT, and HUGO for B) was provided to authors. Data validation was performed by human experts and algorithmically, testing expert-curated relationships for validity in Uberon by querying Ubergraph^128^, a knowledge graph combining mutually referential OBO ontologies including CL and Uberon and featuring precomputed classifications and relationships. This setup makes it possible to feedback data from the ASCT+B tables to correct and expand Uberon and CL but also to align data modelling efforts across ASCT+B tables, Uberon, and CL. Algorithmic testing and validation are ongoing. Most, but not all, ASCT-B table relationships in the initial 11 tables described here do validate.

The version 1.0 tables, also called master tables, are freely available online^129^, and they capture 1,534 AS, 622 CT, 2,154 B (1,492 BG and 632 BP) that are linked by 3,393 AS-AS, 14,987 AS-CT, and 3,580 CT-B relationships supported by 293 unique scholarly publications and 506 web links, see Supplemental Materials.

Like the first maps of our world, the first ASCT+B tables are imperfect and incomplete (see Limitations section). However, they exemplify the utility of ASCT+B tables to standardize and digitize existing data and expertise by pathologists, anatomists, and surgeons at the gross anatomical level; biologists, computer scientists, and others at the single-cell level; and chemists, engineers, and others at the biomarker level.

## 4. 3D Reference Object Library

The spatial location of CT within AS matters as does the context (e.g., the number and type of other CT within the same AS). A 3D reference object library was compiled to capture the size, shape, position, and rotation of major AS in the organ-specific ASCT+B tables. To create reference organs for the 11 initial organs, experts collaborated closely with medical designers to develop anatomically correct, vector-based objects that correctly represent human anatomy, and are properly labelled using the ontology terms captured in the ASCT+B tables.

All 3D reference organs—except for the brain, intestine, and lymph node—use data from the Visible Human Project (VH) male and female dataset made available by the National Library of Medicine^130^. The brain uses the 141 AS of the “Allen Human Reference Atlas – 3D, 2020” representing one half of the human brain^5^. The AS were mirrored to arrive at a whole human brain (as intended) and resized to fit the VH male and female bodies. A 3D model of the male large intestine was kindly provided by Arie Kaufman, Stony Brook University, modified to fit into the VH male body, and used to guide the design of the female large intestine. The lymph node was created using mouse data and the clearing-enhanced 3D method developed by Weizhe Li, Laboratory of Immune System Biology, National Institute of Allergy and Infectious Diseases, NIH^131^. While the size and cellular composition of mouse and human lymph nodes varies, the overall anatomy is well-conserved between species.

The resulting 26 organ objects (male/female and left/right versions exist for kidney and lymph nodes) that are properly positioned in the male and female VH bodies are freely available online^132^. They capture a total of 1,185 unique 3D structures (e.g., the left female kidney has 11 renal papillae; see complete listing at^132^ with 557 unique Uberon terms (including the name of the organ) and Supplemental Materials. Files are provided in GLB format for easy viewing in Babylon.js in a web browser^133^ and in user interfaces (see examples in Tissue Registration and Exploration).

## 5. Reference Atlas Usage

The ASCT+B tables and associated 3D reference organs provide a starting point for building a human reference atlas with common nomenclature for major entities and relationships plus cross-references to existing ontologies and supporting literature. Although the current table format is rather basic, the tables have proven valuable for addressing the originally stated four challenges: (1) to support experimental data registration and annotation across organs and scales (see next subsection); (2) to compare and integrate data from different assay types (see Data Integration and Comparison); (3) to compare healthy and disease data (see Understanding Disease); (4) and to evaluate progress toward the compilation of the human reference atlas (see Measuring Progress).

### Tissue Registration and Exploration

Like any atlas, a human reference atlas is a collection of maps that capture a 3D reality. Like Google maps, the human reference atlas needs to support panning and zoom—from the whole body (macro scale, meters), to the organ level (meso scale, centimeters), to the level of functional tissue units (e.g., alveoli in lung, crypts on colon, glomeruli in kidney, millimeters), down to the single-cell level (micro scale, micrometers). To be usable, the maps in a human reference atlas must use the same index terms and a unifying topological coordinate system so that cells and anatomical structures in adjacent overlapping maps or at different zoom levels can be uniquely named and properly aligned.

The ASCT+B tables and 3D reference organs constitute an agreed-upon framework for experimental data annotation and exploration across organs and scales (i.e., from the entire body down to organs, tissues, cell types, and biomarkers). For example, they are used to support spatially and semantically explicit registration of new tissue data as well as spatial and semantic search, browsing, and exploration of human tissue data. The HuBMAP Registration User Interface (RUI, **Fig. 2a**)^134^ and Exploration User Interface (EUI, **Fig. 2b**)^135^ available via the HuBMAP portal^136^. The code for both user interfaces is freely available on GitHub, and several consortia have registered human tissue samples and are using the exploration user interface to explore their tissue datasets in the context of human anatomy.

**Fig. 2.**
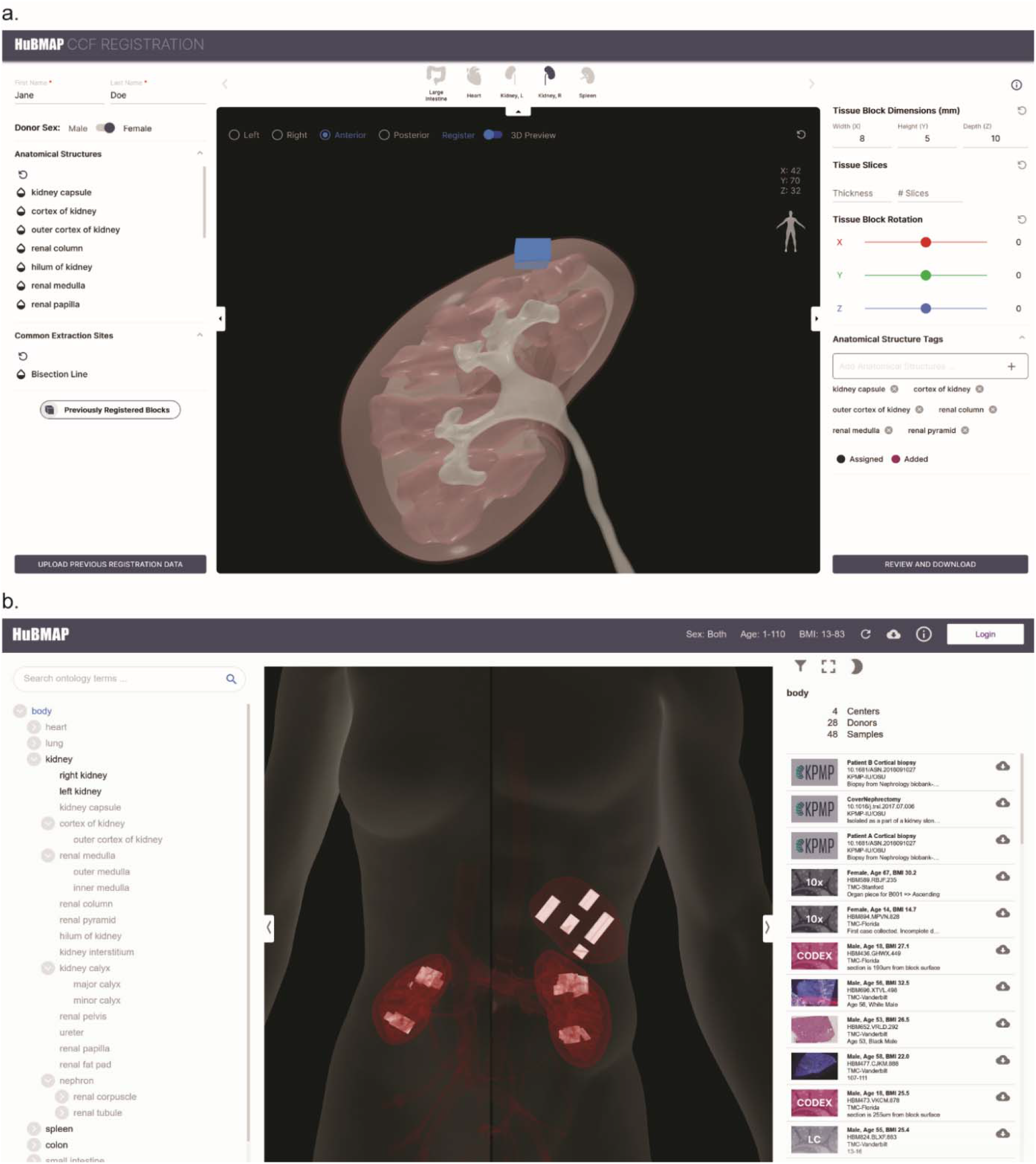
**a**, Registration and semantic annotation of tissue data (blue block) in 3D via collision detection in the Registration User Interface (RUI): A user sizes, positions, and rotates tissue blocks and saves results in JSON format. **b**, Semantically and spatially explicit search, browsing, and filtering of tissue data (white blocks in spleen and kidney) in the Exploration User Interface (EUI): RUI-registered tissue data can be explored semantically using the AS partonomy on left and spatially using the anatomy browser in middle; a filter in top left supports subsetting by sex, age, tissue provider, etc. Clicking on a tissue sample on the right links to the Vitessce image viewer^142^.

### Data Integration: Comparing Cell States Across Different Tissues and in Disease

The ASCT+B framework with controlled ontology vocabulary provides a “lookup table” for AS and their CT composition across organs—for cells formed within and resident in a specific tissue (e.g., epithelia and stroma) as well as cells that migrate across tissues (e.g., immune cells). Immune cells originate primarily in the bone marrow in postnatal life with adaptive lymphocytes subsequently differentiating and maturing in lymphoid tissues such as the thymus and spleen prior to circulating to non-lymphoid tissues and lymph nodes (see **Fig. 3a)**. Therefore, these cells recur across the ASCT+B tables in both the lymphoid (bone marrow, thymus, spleen, lymph node) and non-lymphoid (brain, heart, kidney, lung, skin) tissue tables. Existing data supports a more nuanced and tissue-specific, ontology-based assignment of blood and immune cells. For example, scRNAseq data enables deep phenotyping of hematopoietic stem cells (HSCs) and progenitor cells (HSPCs) and their differentiated progenies across various tissues. When comparing fetal liver *vs*. thymic cell states (**Fig. 3a, right**), a small region of the HSPCs highlighted as “lymphoid progenitors” is shared across the two organs, indicating cells that have migrated from liver to thymus^36, 110^. Data integration defines molecules (e.g., chemokine receptors) that determine tissue residency *versus* migratory properties. These biomarkers (“B”) in turn define tissue-resident vs. migratory cell states, which can be added to the ASCT+B tables to further refine cellular ontology.

**Fig. 3.**
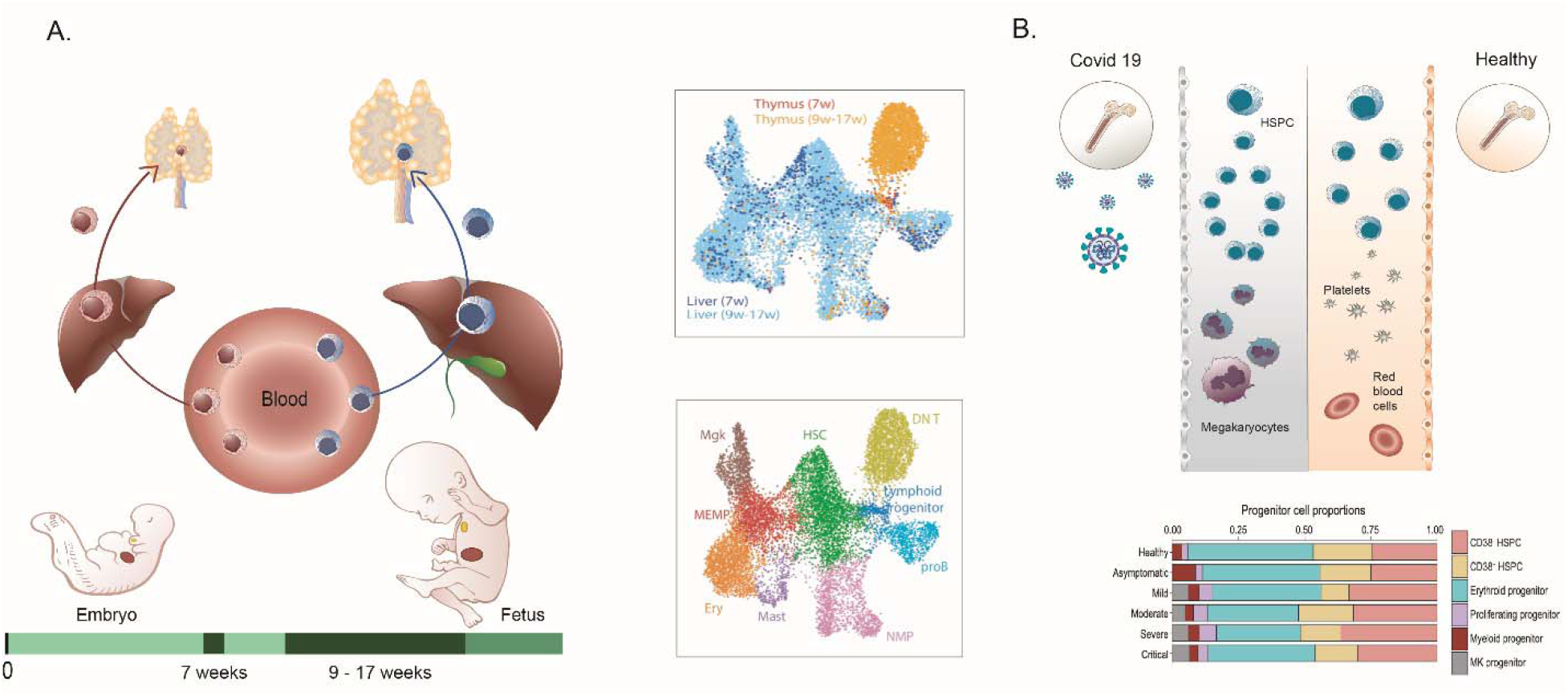
**a**, HSCs migrate from liver to the thymus during embryonic and fetal development. The transcriptomic identity of these HSCs changes between the first and second trimester pregnancy, as shown by the maroon *versus* blue shading of the HSCs in the left (embryo) and right (fetal) part of the figure. The scRNAseq data in the UMAP plots are from^110^; the top plot shows liver cells (blue) and thymus cells (orange) overlapping, labelled “lymphoid progenitor” in bottom plot. **b**, The nature of HSC subsets in adult blood shifts in health *versus* COVID-19. HSCs in the blood of COVID-19 patients (top-left) shows a megakaryocyte priming bias when compared to healthy (top-right). This is quantified in the histogram from^110^ of relative HSPC contributions for different donor/patient cohorts.

A well-annotated healthy human reference atlas can then be used to understand the molecular and cellular alterations in response to perturbations such as infection. Data from several single-cell multi-omics studies of patients’ blood can be federated to compute cellular response during COVID-19 pathogenesis, including HSC progenitor states emerging during disease^137^ (**Fig. 3b)**.

### Understanding Disease

The ASCT+B tables are a semantically and spatially explicit reference for healthy tissue data that can be used to identify changes in molecular states in normal aging or disease. For example, the kidney master table links relevant AS, CT, B to disease and other ontologies for increasing our understanding of disease states. For example, the top significant and specific biomarkers in each reference cell/state cluster might differ in disease or in a cell undergoing repair, regeneration, or in a state of failed or maladaptive repair. Loss of expression or alteration in cellular distribution of an AS specific biomarker may also provide clues to underlying disease. The KPMP is working towards ASCT+B tables that characterize disease by focusing on biomarkers that have important physiological roles in maintaining cellular architecture or function or reveal shifts in cell types associated with acute and chronic diseases. Changes in marker genes in healthy and injured cells provide information on underlying biological pathways and genes that drive these shifts and thus provide critical insights into pathogenetic mechanisms. For example, the *NPHS1* gene, which codes for nephrin in the kidney, is one of the top markers of healthy podocytes and is essential for glomerular function. Mutation in *NPHS1* may be found in patients with proteinuria. The kidney ASCT+B table records that BG *NPHS1* (see lower right in **Fig. 1c**) is expressed in the podocytes of the kidney, and ontology suggests injury to podocytes and glomerular function may cause proteinuria (**Fig. 4**). Ontology IDs provided for AS, CT, B facilitate linkages to clinicopathological knowledge and help provide broader insights into disease^138^. For example, the ASCT+B kidney master table and snRNAseq atlas data^54^ have been used to reveal the cellular identity of diabetic nephropathy genes by distinguishing the healthy interstitium from a diabetic one^139^. Note that some of the existing data are not at the single-cell level; in these cases, regional data (e.g., data bounded by tissue blocks registered within reference organs with known AS, CT, and B—see the RUI and EUI discussion above) can be compared to the kidney master table. In sum, ASCT+B tables interlinked with existing ontologies provide a foundation for new data analysis and the functional study of diseases.

**Fig. 4.**
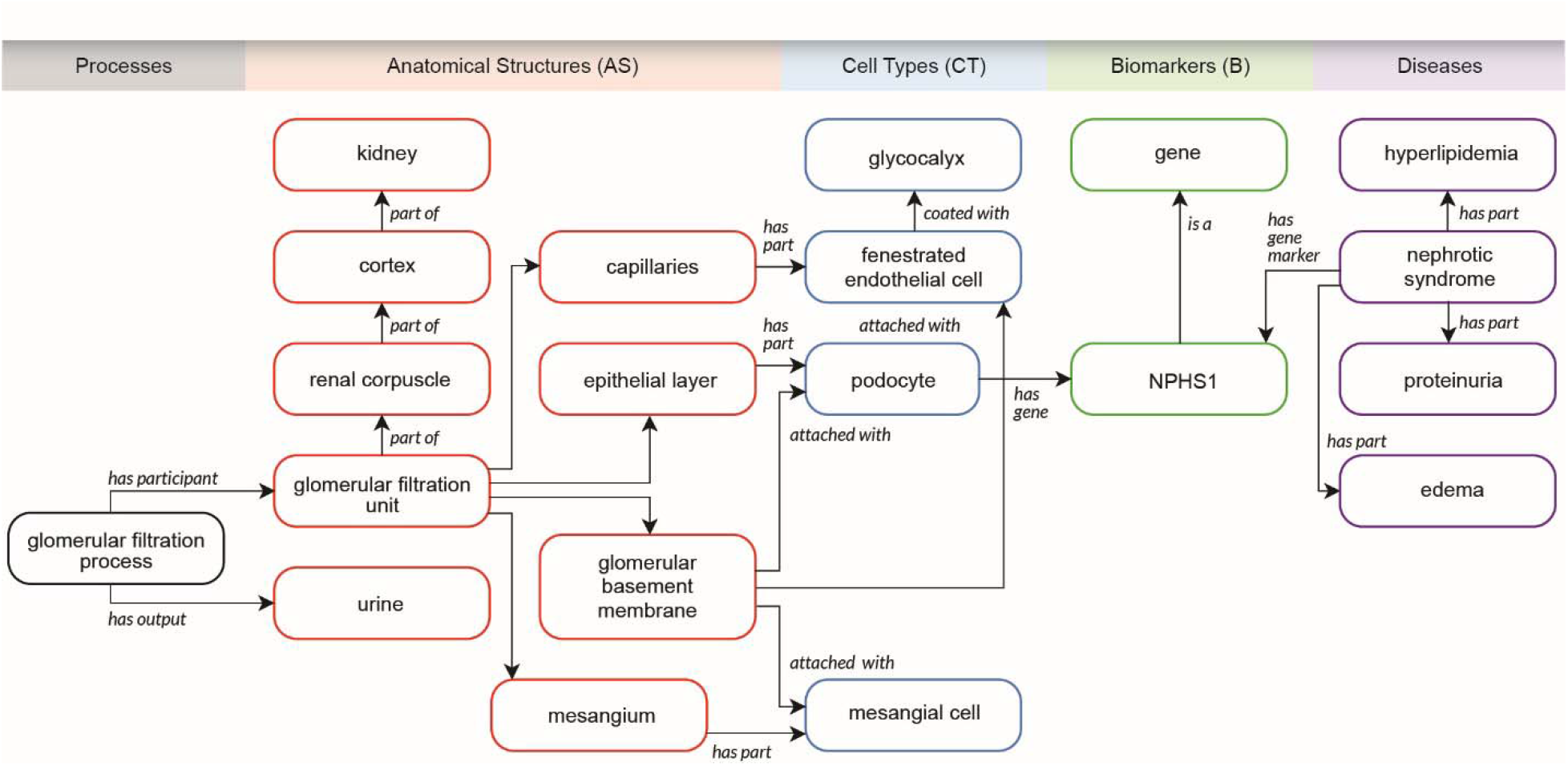
Kidney ASCT+B table linked to processes, disease, clinicopathologic information, and ontology data.

### Measuring Progress

The ASCT+B tables provide a framework to help track progress towards an accurate and complete human reference atlas. Given a scholarly publication that includes a new ASCT+B table, that table can be compared with existing master tables and the number and type of identical (confirmatory) and different (new) AS, CT, and B as well as their relations can be determined. Analogously, the value of a new data release for reference atlas design can be evaluated in terms of the number and type of (new) AS, CT, B and their relations. The ASCT+B Reporter^140^ supports the visual exploration and comparison of ASCT+B tables. Table authors and reviewers can use this online service to upload new tables, examine them visually (see **Fig. 2**), and compare them to existing master tables. When adopted by publishers and editors, the ASCT+B tables provide objective measures and incentives for computing progress towards a human reference atlas. Power analysis methods can be run to assess the coverage and completeness of cell states and/or types and decide what tissues and cells should be sampled next^141^ in support of a data-driven experimental design.

As the number of published and data-derived ASCT+B tables grows, estimates can be run to determine the likely accuracy of AS, CT, B and relations. Entities and relations with much high-quality evidence are more likely to be correct then those with limited or no evidence. Incomplete data can be easily identified and flagged (e.g., AS with no linkages to CT and CT with no B indicate missing data). Given author information for publications and data releases, experts on AS, CT, B, and their relations can be identified and invited to improve tables further.

Using the ASCT+B tables, biases in sampling with respect to donor demographics (e.g., from convenience samples as opposed to using sampling strategies that reflect global demographics), organs (e.g., based on availability of funding), or cell types (e.g., loss of CT due to differential viability or capture efficiency) can be determined and need to be proactively addressed to arrive at an atlas that truly captures healthy human adults.

## 6. Limitations

We designed the format of the ASCT+B tables and 3D reference organs to be easy for experts across many different domains to author and use the tables. They are intended to be intuitive and not require any extensive training in order for them to achieve their intended purpose.

Unfortunately, this means that the current set of tables and reference organs do not capture the complete complexity of the human body. For each organ table, experts recorded the process they used to construct the table, which often included simplifying the anatomy to fit within a strict partonomy, making subjective decisions about which cell types and biomarkers had sufficient evidence to be included in the table, or ignoring normal dynamic changes that occur in the organ over time. For several organs, the B are preliminary and are expected to improve in coverage and robustness in the future.

## 7. Outlook

ASCT+B tables in combination with the 3D reference library provide a unified framework for experimental data annotation and exploration across different levels (i.e., organ, tissue, cell type, and biomarker). The construction and validation of the tables are iterative. Initially, ontology and publication data, along with expertise by organ experts, is codified and unified. Later, experimental datasets will be compared with existing master tables—confirming a subset of all entities and relations captured in it and adding new ones as needed to capture healthy human tissue data. In the near future, the CT typology will be expanded from one level to multiple, making it possible to compare AS partonomy and CT typology datasets at different levels of resolution. New organs will be added to the 3D reference library and micro-anatomical structures such as glomeruli in the kidney, crypts in the large intestine, and alveoli in the lung will be included.

The number of AS, CT, and robust B is likely to increase as new single-cell technologies and computational workflows are developed. Thus, the tables and associated reference objects are a living “snapshot” of the status of the collective work toward a human reference atlas, against which experimentalists can calibrate their data and ultimately contribute to the atlas by expanding or refining it. Future uses of the human reference atlas might include cross-species comparisons or cross-species annotations, cross-tissue/organ comparisons, comparisons of healthy versus common or rare genetic variations, and usage in teaching—expanding widely used anatomy books^62, 120^ to the single-cell level.

It will take much effort and expertise to arrive at a consensus human reference atlas and to develop methods and user interfaces that utilize it to advance research and improve human health. Experts interested in contributing to this international and interdisciplinary effort are invited to register via https://iu.co1.qualtrics.com/jfe/form/SV_bpaBhIr8XfdiNRH to receive more information and meeting invites.

## Acknowledgements

We are grateful to Blue Lake from University of California, San Diego, for assistance with annotations and analyzing the snRNAseq HUBMAP data for several of the markers in the kidney ASCT+B tables, Becky Steck and Rachel Dull from University of Michigan for assistance with the nomenclature and curation of kidney partonomy, and Seth Winfree, IUPUI for valuable discussions regarding the kidney ASCT+B table. We thank the Kidney Precision Medicine Project (KPMP), especially the Tissue Interrogation Sites and the Controlled Cell Vocabulary working group for guidance and development of the initial sets of ASCT+B kidney tables. We acknowledge Li Yao for segmenting and optimizing the mouse popliteal lymph node model from high-resolution microscopic data. The work was funded, in part, by NIH Awards OT2OD026671, U54DK120058, 1UH3CA246594, 1U54AI142766, 1UG3CA256960, 1UG3HL145609, U54HL145608, U54HL145611, UH3DK114933, DK110814 and DK107350; National Institute of Allergy and Infectious Diseases (NIAID), Department of Health and Human Services under BCBB Support Services Contract HHSN316201300006W/HHSN27200002; the Intramural Research Program of the NIH at NIAID; and Helmsley Charitable Trust 2018PG-T1D071.

## Supplemental Information

### Contents

S1. ASCT+B Tables

S2. CCF 3D Reference Object Library

## S1. ASCT+B Tables

Anatomical Structures, Cell Types, plus Biomarkers (ASCT+B) tables aim to capture the structure of anatomical human body parts, the typology of cells, and biomarkers used to identify cell types (e.g., gene and protein). The tables are authored and reviewed by an international team of anatomists, pathologists, physicians, and other experts.

The v1.0 release features a total of 11 tables.

| Table Name | DOI |
| --- | --- |
| 1. Bone Marrow & Blood/Pelvis v1.0 | https://doi.org/10.48539/HBM963.TBFP.428 |
| 2. Brain v1.0 | https://doi.org/10.48539/HBM264.CCRF.525 |
| 3. Heart v1.0 | https://doi.org/10.48539/HBM873.NNVD.559 |
| 4. Intestine, Large v1.0 | https://doi.org/10.48539/HBM287.KVCD.758 |
| 5. Kidney v1.0 | https://doi.org/10.48539/HBM325.PTQS.258 |
| 6. Lung v1.0 | https://doi.org/10.48539/HBM868.DWJZ.874 |
| 7. Lymph Node v1.0 | https://doi.org/10.48539/HBM375.LMPC.837 |
| 8. Skin v1.0 | https://doi.org/10.48539/HBM227.WHPG.367 |
| 9. Spleen v1.0 | https://doi.org/10.48539/HBM625.VPLJ.455 |
| 10. Thymus v1.0 | https://doi.org/10.48539/HBM934.JMVN.957 |
| 11. Vasculature v1.0 | https://doi.org/10.48539/HBM426.GZPN.888 |

## S2. CCF 3D Reference Object Library

The CCF 3D Reference Object Library comprises anatomically correct reference organs. The organs are developed by specialists in 3D medical illustration and approved by organ experts.

The v1.0 release features a total of 26 reference organs.

The crosswalk table to the ASCT+B tables is available via the CCF 3D Reference Object Library page at https://hubmapconsortium.github.io/ccf/pages/ccf-3d-reference-library.html.

| 3D Reference Organ Name | DOI |
| --- | --- |
| 1. Pelvis, Male v1.0 | https://doi.org/10.48539/HBM672.KSST.448 |
| 2. Pelvis, Female v1.0 | https://doi.org/10.48539/HBM539.RRDB.894 |
| 3. Brain, Male v1.0 | https://doi.org/10.48539/HBM927.XKJR.426 |
| 4. Brain, Female v1.0 | https://doi.org/10.48539/HBM438.XKJT.666 |
| 5. Heart, Male v1.0 | https://doi.org/10.48539/HBM384.FDPD.646 |
| 6. Heart, Female v1.0 | https://doi.org/10.48539/HBM894.FGQN.237 |
| 7. Intestine, Large, Male v1.0 | https://doi.org/10.48539/HBM235.QFTP.824 |
| 8. Intestine, Large, Female v1.0 | https://doi.org/10.48539/HBM734.KXRL.243 |
| 9. Kidney, Male, Left v1.0 | https://doi.org/10.48539/HBM792.NPXX.335 |
| 10. Kidney, Male, Right v1.0 | https://doi.org/10.48539/HBM693.RTRN.866 |
| 11. Kidney, Female, Left v1.0 | https://doi.org/10.48539/HBM765.JWFB.892 |
| 12. Kidney, Female, Right v1.0 | https://doi.org/10.48539/HBM726.HWKT.534 |
| 13. Lung, Male v1.0 | https://doi.org/10.48539/HBM373.VPSC.692 |
| 14. Lung, Female v1.0 | https://doi.org/10.48539/HBM467.NHKR.974 |
| 15. Lymph Node, Male, Left v1.0 | https://doi.org/10.48539/HBM668.RBLW.256 |
| 16. Lymph Node, Male, Right v1.0 | https://doi.org/10.48539/HBM699.BLKS.298 |
| 17. Lymph Node, Female, Left v1.0 | https://doi.org/10.48539/HBM897.HVMV.989 |
| 18. Lymph Node, Female, Right v1.0 | https://doi.org/10.48539/HBM439.ZTGH.778 |
| 19. Skin, Male v1.0 | https://doi.org/10.48539/HBM689.CGPV.442 |
| 20. Skin, Female v1.0 | https://doi.org/10.48539/HBM287.TJZD.872 |
| 21. Spleen, Male v1.0 | https://doi.org/10.48539/HBM583.DRHV.265 |
| 22. Spleen, Female v1.0 | https://doi.org/10.48539/HBM626.WZGS.532 |
| 23. Thymus, Male v1.0 | https://doi.org/10.48539/HBM955.BGTP.947 |
| 24. Thymus, Female v1.0 | https://doi.org/10.48539/HBM896.TWPT.542 |
| 25. Vasculature, Male v1.0 | https://doi.org/10.48539/HBM784.ZDZF.357 |
| 26. Vasculature, Female v1.0 | https://doi.org/10.48539/HBM253.DKWB.737 |

## References

1. International Human Genome Sequencing (IHGS) Consortium Initial sequencing and analysis of the human genome. Nature 409, 860–921 (2001).

2. Venter, J.C. et al. The sequence of the human genome. Science 291, 1304–1351 (2001).

3. Regev, A. et al. The Human Cell Atlas. eLife 6, e27041 (2017).

4. Rozenblatt-Rosen, O., Stubbington, M.J.T., Regev, A. & Teichmann, S.A. The Human Cell Atlas: from vision to reality. Nature 574, 187–192 (2017).

5. Ding, S.L. et al. Comprehensive cellular-resolution atlas of the adult human brain. J Comp Neurol 524, 3127–3481 (2016).

6. Devor, A. et al. The challenge of connecting the dots in the B.R.A.I.N. Neuron 80, 270– 274 (2013).

7. Moghe, I., Loupy, A. & Solez, K. The Human Cell Atlas Project by the numbers: relationship to the Banff Classification. Am J Transplant 18, 1830.

8. Lonsdale, J. et al. The Genotype-Tissue Expression (GTEx) project. Nat Genet 45, 580– 585 (2013).

9. McMahon, A.P. et al. GUDMAP: the genitourinary developmental molecular anatomy project. J Am Soc Nephrol 19, 667–671 (2008).

10. Srivastava, S. et al. The making of a PreCancer Atlas: promises, challenges, and opportunities. Trends Cancer 4, 523–536 (2018).

11. Snyder, M.P. et al. The human body at cellular resolution: the NIH Human Biomolecular Atlas Program. Nature 574, 187–192 (2019).

12. El-Achkar, T.M. et al. A multimodal and integrated approach to interrogate human kidney biopsies with rigor and reproducibility: guidelines from the Kidney Precision Medicine Project. Physiol Genomics 53, 1–11 (2021).

13. Ardini-Poleske, M.E. et al. LungMAP: the Molecular Atlas of Lung Development Program. Am J Physiol Lung Cell Mol Physiol 313, L733–L740 (2013).

14. United Kingdom Research and Innovation (UKRI) Medical Research Council. https://mrc.ukri.org (2021).

15. Oxburgh, L. et al. (Re)building a kidney. J Am Soc Nephrol 28, 1370 (2017).

16. NIH Stimulating Peripheral Activity to Relieve Conditions (SPARC). https://commonfund.nih.gov/sparc (2020).

17. Heng, H.H.Q. Cancer genome sequencing the challenges ahead. Bioessays 29, 783–794 (2007).

18. TGCA Research Network Comprehensive genomic characterization defines human glioblastoma genes and core pathways. Nature 455, 1061–1068 (2008).

19. Freire, P. et al. Exploratory analysis of the copy number alterations in glioblastoma multiforme. PLOS ONE 3, e4076 (2008).

20. Miller, J.A. et al. Common cell type nomenclature for the mammalian brain. eLife 9, e59928 (2020).

21. Ding, S.L. et al. Allen Human Reference Atlas – 3D, 2020 http://download.alleninstitute.org/informatics-archive/allen_human_reference_atlas_3d_2020/version_1 (2020)

22. Fonseca, C.G. et al. The Cardiac Atlas Project: an imaging database for computational modeling and statistical atlases of the heart. Bioinformatics 27, 2288–2295 (2011).

23. Géron, A., Werner, J., Wattiez, R., Lebaron, P. & Mattallana-Surget, S. Deciphering the functioning of microbial communities: shedding light on the critical steps in metaproteomics. Front Microbiol 10, 2395 (2019).

24. Manz, M.G., Miyamoto, T., Akashi, K. & Weissman, I.L. Prospective isolation of human clonogenic common myeloid progenitors. Proc Natl Acad Sci U S A 99, 11872–11877 (2002).

25. Fajtova, M., Kovarikova, A., Svec, P., Kankuri, E. & Sedlak, J. Immunophenotypic profile of nucleated erythroid progenitors during maturation in regenerating bone marrow. Leuk Lymphoma 54, 2523–2530 (2013).

26. Kawamura, S. et al. Identification of a human clonogenic progenitor with strict monocyte differentiation potential: a counterpart of mouse cMoPs. Immunity 46, 835–848 (2017).

27. Mousset, C.M. et al. Comprehensive phenotyping of t cells using flow cytometry. Cytometry A 95, 647–654 (2019).

28. Mello, F.V. et al. Maturation-associated gene expression profiles along normal human bone marrow monopoiesis. Br J Haematol 176, 464–474 (2017).

29. Tomer, A. Human marrow megakaryocyte differentiation: multiparameter correlative analysis identifies von Willebrand factor as a sensitive and distinctive marker for early (2N and 4N) megakaryocytes. Blood 104, 2722–2727 (2004).

30. Doulatov, S. et al. Revised map of the human progenitor hierarchy shows the origin of macrophages and dendritic cells in early lymphoid development. Nat Immunol 11, 585–593 (2010).

31. Elghetany, M.T., Ge, Y., Patel, J., Martinez, J. & Uhrova, H. Flow cytometric study of neutrophilic granulopoiesis in normal bone marrow using an expanded panel of antibodies: correlation with morphologic assessments. J Clin Lab Anal 18, 36–41 (2004).

32. Szabo, P.A. et al. Single-cell transcriptomics of human T cells reveals tissue and activation signatures in health and disease. Nat Commun 10, 4706 (2019).

33. Kaminski, D.A., Wei, C., Qian, Y., Rosenberg, A.F. & Sanz, I. Advances in human B cell phenotypic profiling. Front Immunol 3, 302 (2012).

34. Hay, S.B., Ferchen, K., Chetal, K., Grimes, H.L. & Salomonis, N. The Human Cell Atlas bone marrow single-cel interactive web portal. Exp Hematol 68, P51–61 (2018).

35. Clavarino, G. et al. Novel strategy for phenotypic characterization of human B lymphocytes from precursors to effector cells by flow cytometry. PLOS ONE 11, e0162209 (2016).

36. Popescu, D.M. et al. Decoding human fetal liver haematopoiesis. Nature 574, 365–371 (2019).

37. Hodge, R.D. et al. Conserved cell types with divergent features in human versus mouse cortex. Nature 573, 61–68 (2019).

38. Hawrylycz, M.J. et al. An anatomically comprehensive atlas of the adult human brain transcriptome. Nature 489, 391–399 (2012).

39. Litvinuková, M. et al. Cells of the adult human heart. Nature 588, 466–472 (2020).

40. Tucker, N.R. et al. Transcriptional and cellular diversity of the human heart. Circulation 142, 466–482 (2020).

41. Giannasca, P.J., Giannasca, K.T., Leichtner, A.M. & Neutra, M.R. Human intestinal M cells display the sialyl Lewis A antigen. Infect Immun 67, 946–953 (1999).

42. Buettner, M. & Lochner, M. Development and function of secondary and tertiary lymphoid organs in the small intestine and the colon. Front Immunol 7, 342 (2016).

43. Hoyle, C.H. & Burnstock, G. Neuronal populations in the submucous plexus of the human colon. J Anat 166, 7–22 (1989).

44. Westerhoff, M. & Greeson, J. Colon, in Histology for Pathologists, Edn. 5th. (ed. S. Mills) 640–663 (Wolters Kluwer, Philadelphia; 2019).

45. Azzali, G. Structure, ymphatic vascularization and lymphocyte migration in mucosa-associated lymphoid tissue. Immunol Rev 195, 178–189 (2003).

46. Furness, J.B., Callaghan, B.P., Rivera, L.R. & Cho, H.J. The enteric nervous system and gastrointestinal innervation: integrated local and central control. Adv Exp Med Biol 817, 39–71 (2014).

47. Arai, T. & Kino, I. Morphometrical and cell kinetic studies of normal human colorectal mucosa. Comparison between the proximal and the distal large intestine. Acta Pathol Jpn 39, 725–730 (1989).

48. Fenton, T.M. et al. Immune Profiling of Human Gut-Associated Lymphoid Tissue Identifies a Role for Isolated Lymphoid Follicles in Priming of Region-Specific Immunity. Immunity 52, 557–570 (2020).

49. Habowski, A.N. et al. Transcriptomic and proteomic signatures of stemness and differentiation in the colon crypt. Commun Biol 3, 453 (2020).

50. Lundqvist, C., Baranov, V., Hammarström, S., L Athlin & Hammarström, M.L. Intra-epithelial lymphocytes. Evidence for regional specialization and extrathymic T cell maturation in the human gut epithelium. Int Immunol 7, 1473–1487 (1995).

51. Lockyer, M.G. & Petras, R.E. Appendix, in Histology for Pathologists. (ed. S. Mills) 664– 676 (Wolters Kluwer, Philadelphia; 2019).

52. Pittman, M.E. & Yantiss, R.K. Anal canal, in Histology for Pathologists. (ed. S. Mills) 677–691 (Wolters Kluwer, Philadelphia; 2019).

53. Kriz, W. & Bankir, L. A standard nomenclature for structures of the kidney. The Renal Commission of the International Union of Physiological Sciences (IUPS). Kidney Int 33, 1–7 (1988).

54. Lake, B.B. et al. A single-nucleus RNA-sequencing pipeline to decipher the molecular anatomy and pathophysiology of human kidneys. Nat Commun 10, 2832 (2019).

55. Barry, D.M. et al. Molecular determinants of nephron vascular specialization in the kidney. Nat Commun 10, 5705 (2019).

56. Menon, R. et al. Single cell transcriptomics identifies focal segmental glomerulosclerosis remission endothelial biomarker. JCI Insight 5, e133267 (2020).

57. Ransick, A. et al. Single-cell profiling reveals sex, lineage, and regional diversity in the mouse kidney. Dev Cell 51, 399–413 (2019).

58. Limbutara, K., Chou, C.L. & Knepper, M.A. Quantitative proteomics of all 14 renal tubule segments in rat. J Am Soc Nephrol 31, 1255–1266 (2020).

59. Kuppe, C. et al. Decoding myofibroblast origins in human kidney fibrosis. Nature 589, 281–286 (2021).

60. Stewart, B.J. et al. Spatiotemporal immune zonation of the human kidney. Science 365, 1461–1466 (2019).

61. Kirita, Y., Wu, H., Uchimura, K., Wilson, P.C. & Humphreys, B.D. Cell profiling of mouse acute kidney injury reveals conserved cellular responses to injury. Proc Natl Acad Sci U S A 117, 15874–15883 (2020).

62. Standring, S. Gray’s Anatomy: The Anatomical Basis of Clinical Practice. (Elsevier, Edinburgh; 2016).

63. Haefeli-Bleuer, B. & Weibel, E.R. Morphometry of the human pulmonary acinus. Anat Rec 220, 401–414 (1988).

64. Whitsett, J.A., Kalin, T.V., Xu, Y. & Kalinichenko, V.V. Building and regenerating the lung cell by cell. Physiol Rev 99, 513–554 (2019).

65. Plasschaert, L.W. et al. A single-cell atlas of the airway epithelium reveals the CFTR-rich pulmonary ionocyte. Nature 560, 377–381 (2018).

66. Xu, Y. et al. Single-cell RNA sequencing identifies diverse roles of epithelial cells in idiopathic pulmonary fibrosis. JCI Insight 1, e90558 (2016).

67. Adams, T.S. et al. Single-cell RNA-seq reveals ectopic and aberrant lung-resident cell populations in idiopathic pulmonary fibrosis. Sci Adv 6, eaba1983 (2020).

68. Wang, A. et al. Single-cell multiomic profiling of human lungs reveals cell-type-specific and age-dynamic control of SARS-CoV2 host genes. eLife 9, e62522 (2020).

69. Travaglini, K.J. et al. A molecular cell atlas of the human lung from single-cell RNA sequencing. Nature 587, 619–625 (2020).

70. Deprez, M. et al. A single-cell atlas of the human healthy airways. Am J Respir Crit Care Med 202, 1636–1645 (2019).

71. Sun, X. & Morrisey, E. Lung cell census: a view from LungMAP. (In Preparation).

72. Medeiros, L.J. et al. Normal anatomy and function of lymph nodes and spleen, in Tumors of the Lymph Node and Spleen (American Registry of Pathology, Silver Spring, MD; 2017).

73. Medeiros, L.J. et al. Tumors of the Lymph Node and Spleen. (American Registry of Pathology, Silver Spring, MD; 2017).

74. O’Malley, D.P., George, T.I., Orazi, A. & Abbondanzo, S.L. General immunology of lymph node and spleen, in Benign and Reactive Conditions of Lymph Node and Spleen (American Registry of Pathology, Silver Spring, MD; 2009).

75. Angel, C.E. et al. Distinctive localization of antigen-presenting cells in human lymph nodes. Blood 113, 1257–1267 (2009).

76. James, K.R. et al. Distinct microbial and immune niches of the human colon. Nat Immunol 21, 343–353 (2020).

77. Link, A. et al. Association of T-zone reticular networks and conduits with ectopic lymphoid tissues in mice and humans. Am J Pathol 178, 1662–1675 (2011).

78. Park, S.M. et al. Mapping the distinctive populations of lymphatic endothelial cells in different zones of human lymph nodes. PLOS ONE 9, e106814 (2014).

79. Xiang, M. et al. A single-cell transcriptional roadmap of the mouse and human lymph node lymphatic vasculature. Front Cardiovasc Med 7, 52 (2020).

80. Takeda, A. et al. Single-cell survey of human lymphatics unveils marked endothelial cell heterogeneity and mechanisms of homing for neutrophils. Immunity 51, 561–572 (2019).

81. Kunicki, M.A., Hernandez, L.C.A., Davis, K.L., Bacchetta, R. & Roncarolo, M.G. Identity and diversity of human peripheral Th and T regulatory cells defined by single-cell mass cytometry. J Immunol 200, 336–346 (2018).

82. Pusztaszeri, M.P., Seelentag, W. & Bosman, F.T. Immunohistochemical expression of endothelial markers CD31, CD34, von Willebrand factor, and Fli-1 in normal human tissues. J Histochem Cytochem 54, 385–395 (2006).

83. Reynolds, G. et al. Developmental cell programs are co-opted in inflammatory skin disease. Science 371, 364 (2021).

84. Dyring-Anderson, B. et al. Spatially and cell-type resolved quantitative proteomic atlas of healthy human skin. Nat Commun 11, 5587 (2020).

85. Fuchs, E. Keratins and the skin. Annu Rev Cell Dev Biol 11, 123–153 (1995).

86. Nestle, F.O., Meglio, P.D., Qin, J.Z. & Nickoloff, B.J. Skin immune sentinels in health and disease. Nat Rev Immunol 9, 679–691 (2009).

87. Laverdet, B. et al. Skin innervation: important roles during normal and pathological cutaneous repair. Histol Hitopathol 30, 875–892 (2015).

88. Ryan, T.J. The blood vessels of the skin. J Invest Dermatol 67, 110–118 (1976).

89. Popescu, D.M. A single cell atlas of adult healthy, psoriatic and atopic dermatitis skin. https://developmentcellatlas.ncl.ac.uk/datasets/hca_skin_portal (2021).

90. Bos, J.D. et al. The skin immune system (SIS): distribution and immunophenotype of lymphocyte subpopulations in normal human skin. J Invest Dermatol 88, 569–573 (1987).

91. Schweizer, J. et al. New consensus nomenclature for mammalian keratins. J Cell Biol 174, 169–174 (2006).

92. Eberl, G., Colonna, M., Di Santo, J.P. & McKenzie, A.N.J. Innate lymphoid cells: a new paradigm in immunology. Science 348, aaa6566 (2015).

93. Ali, N. & Rosenblum, M.D. Regulatory T cells in skin. Immunology 152, 372–381 (2017).

94. Huber, W.E. et al. A tissue-restricted cAMP transcriptional response: SOX10 modulates alpha-melanocyte-stimulating hormone-triggered expression of microphthalmia-associated transcription factor in melanocytes. J Biol Chem 278, 45224–45230 (2003).

95. Haniffa, M., Gunawan, M. & Jardine, L. Human skin dendritic cells in health and disease. J Dermatol Sci 77, 85–92 (2015).

96. Cesta, M.F. Normal structure, function, and histology of the spleen. Toxicol Pathol 34, 455–465 (2006).

97. Madissoon, E. et al. scRNA-seq assessment of the human lung, spleen, and esophagus tissue stability after cold preservation. Genome Biol 21, 1 (2019).

98. Pack, M. et al. DEC-205/CD205+ dendritic cells are abundant in the white pulp of the human spleen, including the border region between the red and white pulp. Immunology 123, 438–446 (2008).

99. Steiniger, B.S. Human spleen microanatomy: why mice do not suffice. Immunology 145, 334–346 (2015).

100. Steiniger, B.S., Seiler, A., Lampp, K., Wilhelmi, V. & Stachniss, V. Blymphocyte compartments in the human splenic red pulp: capillary sheaths and periarteriolar regions. Histochem Cell Biol 141, 507–518 (2014).

101. Steiniger, B.S., Stachniss, V., Schwarzbach, H. & Barth, P.J. Phenotypic differences between red pulp capillary and sinusoidal endothelia help localizing the open splenic circulation in humans. Histochem Cell Biol 128, 391–398 (2007).

102. Qiu, J. et al. The characteristics of vessel lining cells in normal spleens and their role in the pathobiology of myelofibrosis. Blood Adv 2, 1130–1145 (2018).

103. Lewis, S.M., Williams, A. & Eisenbarth, S.C. Structure and function of the immune system in the spleen. Sci Immunol 4, eaau6085 (2019).

104. Mittag, D. et al. Human dendritic cell subsets from spleen and blood are similar in phenotype and function but modified by donor health status. J Immunol 186, 6207–6217 (2011).

105. Cheng, H.W. et al. Origin and differentiation trajectories of fibroblastic reticular cells in the splenic white pulp. Nat Commun 10, 1739 (2019).

106. Van Krieken, J.H.J.M., Te Velde, J., Leenheers-Binnendijk, L. & Van de Velde, C.J.H. The human spleen; a histological study in splenectomy specimens embedded in methylmethacrylate. Histopathology 9, 571–585 (1985).

107. Van Krieken, J.H.J.M. & Te Velde, J. Immunohistology of the human spleen: an inventory of the localization of lymphocyte subpopulations. Histopathology 10, 285–294 (1986).

108. Bautista, J.L. et al. Single-cell transcriptional profiling of human thymic stroma uncovers novel cellular heterogeneity in the thymic medulla. Nat Commun 12, 1096 (2021).

109. Haynes, B.F. The human thymic microenvironment. Adv Immunol 36, 87–142 (1984).

110. Park, J.E. et al. A cell atlas of human thymic development defines T cell repertoire formation. Science 367, eaay3224 (2020).

111. Pearse, G. Normal structure, function and histology of the thymus. Toxicol Pathol 34, 504– 514 (2006).

112. Suster, S. & Rosai, J. Histology of the normal thymus. Am J Surg Pathol 14, 284–303 (1990).

113. Mignini, F. et al. Neuro-immune modulation of the thymus microenvironment (review). Int J Mol Med 33, 1392–1400 (2014).

114. Stoeckle, C. et al. Isolation of myeloid dendritic cells and epithelial cells from human thymus. J Vis Exp 79, e50951 (2013).

115. Marcovecchio, G.E. et al. Thymic epithelium abnormalities in DiGeorge and Down syndrome patients contribute to dysregulation in T cell development. Front Immunol 10, 447 (2019).

116. Wakimoto, T. et al. Identification and characterization of human thymic cortical dendritic macrophages that may act as professional scavengers of apoptotic thymocytes. Immunobiology 213, 837–847 (2008).

117. Lavaert, M. et al. Integrated scRNA-Seq identifies human postnatal thymus seeding progenitors and regulatory dynamics of differentiating immature thymocytes. Immunity 52, 1088–1104 (2020).

118. Nuñez, S. et al. The human thymus perivascular space is a functional niche for viral-specific plasma cells. Sci Immunol 1, eaah4447 (2016).

119. Bendriss-Vermare, N. et al. Human thymus contains IFN-alpha-producing CD11c(-), myeloid CD11c(+), and mature interdigitating dendritic cells. J Clin Invest 107, 835–844 (2001).

120. Netter, F.H. Atlas of Human Anatomy, Edn. 7th. (Elsevier, Philadelphia; 2019).

121. Kandathil, A. & Chamarthy, M. Pulmonary vascular anatomy & anatomical variants. Cardiovasc Diagn Ther 8, 201–207 (2018).

122. Perlmutter, D. & Rhoton Jr, A.L. Microsurgical anatomy of the distal anterior cerebral artery. J Neurosurg 49, 204–228 (1978).

123. Hacein-Bey, L. et al. The ascending pharyngeal artery: branches, anastomoses, and clinical significance. AJNR Am J Neuroradiol 23, 1246–1256 (2002).

124. Vuong, S.M., Jeong, W.J., Morales, H. & Abruzzo, T.A. Vascular diseases of the spinal cord: infarction, hemorrhage, and venous congestive myelopathy. Semin Ultrasound CT MR 37 (2016).

125. Picel, A.C., Hsieh, T.C., Shapiro, R.M., Vezeridis, A.M. & Isaacson, A.J. Prostatic artery embolization for benign prostatic hyperplasia: patient evaluation, anatomy, and technique for successful treatment. Radiographics 39, 1526–1548 (2019).

126. Vummidi, D. et al. Pseudolesion in segment IV A of the liver from vein of Sappey secondary to SVC obstruction. Radiol Case Rep 5, 394 (2015).

127. Osumi-Sutherland, D., Keays, M., Lein, E.S. & Teichmann, S.A. The Human Cell Atlas: cell types and ontologies. Nat Cell Biol (In Press).

128. Balhoff, J. & Curtis, C.K. Ubergraph. https://github.com/INCATools/ubergraph (2021).

129. HuBMAP Consortium CCF Anatomical Structures, Cell Types and Biomarkers (ASCT+B) Tables https://hubmapconsortium.github.io/ccf/pages/ccf-anatomical-structures.html (2021).

130. NLM Visible Human Project (VHP) data sets. https://www.nlm.nih.gov/databases/download/vhp.html (2020).

131. Li, W., Germain, R.N. & Gerner, M.Y. Multiplex, quantitative cellular analysis in large tissue volumes with clearing-enhanced 3D microscopy (Ce3D). Proc Natl Acad Sci U S A 114, E7321–E7330 (2017).

132. HuBMAP Consortium CCF 3D Reference Object Library. https://hubmapconsortium.github.io/ccf/pages/ccf-3d-reference-library.html (2021).

133. Babylon.js Sandbox. https://sandbox.babylonjs.com/ (2021).

134. HuBMAP Consortium CCF Registration User Interface. https://hubmapconsortium.github.io/ccf-ui/rui (2021).

135. HuBMAP Consortium CCF Exploration User Interface. https://portal.hubmapconsortium.org/ccf-eui (2021).

136. NIH Human Biomolecular Atlas Program (HuBMAP) data portal. https://portal.hubmapconsortium.org (2020).

137. Stephenson, E. et al. The immune response in COVID-19 detailed by single cell multiomics. Nat Med (In Press).

138. Ong, E. et al. Modelling kidney disease using ontology: insights from the Kidney Precision Medicine Project. Nat Rev Nephrol 16, 686–696 (2020).

139. Barwinska, D. et al. Molecular characterization of the human kidney interstitium in health and disease. Sci Adv 7, eabd3359 (2021).

140. HuBMAP Consortium ASCT+B Reporter. https://hubmapconsortium.github.io/ccf-asct-reporter (2021).

141. The HCA Consortium The Human Cell Atlas https://www.humancellatlas.org/wp-content/uploads/2019/11/HCA_WhitePaper_18Oct2017-copyright.pdf (2017)

142. Manz, T. et al. Viv: multiscale visualization of high-resolution multiplexed bioimaging data on the web. Preprint at https://doi.org/10.31219/osf.io/wd31212gu (2020).

